# Genotype-by-environment interactions for seminal fluid expression and sperm competitive ability

**DOI:** 10.1101/727685

**Authors:** Bahar Patlar, Steven A. Ramm

## Abstract

Sperm competition commonly occurs whenever females mate multiply, leading to variation in male paternity success. This can be due to variation in the various traits that might affect sperm competitive ability, which itself depends on both genetic and environmental factors, as well as on genotype-by-environment interactions (GEI). Seminal fluid is a major component of the male ejaculate that is often expected to mediate sperm competition, where different genotypes can differ in their seminal fluid expression as a response to different level of sperm competition (i.e., exhibit GEI). We therefore here focussed on testing for GEI in expression of two recently identified seminal fluid transcripts, *suckless-1* and *suckless-2*, which potentially modulate sperm competitive ability in the simultaneously hermaphroditic flatworm *Macrostomum lignano* via their effects on manipulating post-mating partner behaviour and ultimately the fate of transferred ejaculates. In addition, we sought to test for GEI in sperm competitive ability, to investigate the relationship between natural variation in the expression of these seminal fluid transcripts generated through GEI and relative paternity success. To do so, we manipulated social group size, which has been shown to successfully alter sperm competition level in *M. lignano*, in a set of inbred lines (genotypes) and then measured both the expression level of *suckless-1* and *suckless-2* in focal worms together with their relative paternity success in a standardised sperm competition (*P*_1_ & *P*_2_) assay. We found GEI for the expression level of *suckless-1* and *suckless-2*, as well as for sperm competitive ability. Moreover, we found a positive relation between the expression of *suckless-1* and paternity success. This suggests that natural variation in the expression of this seminal fluid transcript indeed can influence sperm competition outcomes in *M. lignano*.

## Introduction

Sperm competition, that is the competition between the ejaculates of two or more males for the fertilization of a given set of ova (Parker, 1970), and cryptic female choice, in which females influence the outcome of sperm competition (Eberhard, 1996), are important evolutionary forces across a diverse range of taxa in which females mate multiply. Variation in paternity success is therefore determined by many factors affecting sperm competitive ability of the ejaculate (Parker, 1970; Lewis & Austad, 1990; Radwan, 1996; Birkhead & Møller, 1998; Gage & Morrow, 2003; Snook, 2005; Simmons & Parker, 2006; Pizzari & Parker, 2009). Numerous adaptations related to the amount and quality of sperm such as sperm number, size, velocity, mobility, and storage capacity affect relative paternity success under sperm competition (Birkhead & Møller, 1998; Wedell *et al.*, 2002; Snook, 2005; Bjork & Pitnick, 2006; Pitnick *et al.*, 2009; Pizzari & Parker, 2009; Parker & Pizzari, 2010; Godwin *et al.*, 2017). Alongside sperm, males typically also transfer large number of seminal fluid proteins/peptides (SFPs) during mating (reviewed by Poiani, 2006; Avila *et al.*, 2011; Hopkins *et al.*, 2017) and these too are theoretically expected to be strongly shaped by sperm competition, favouring the evolution of SFP functions that confer a male fitness benefit through increased competitive fertilization success (Cameron *et al.*, 2007; Dhole & Servedio, 2014; Dapper & Wade, 2016).

SFPs can either decrease the risk of sperm competition, for example by preventing female remating, or increase the chance of sperm to fertilize eggs by serving as offensive or defensive tools against rival sperm in female genital tracts. Their well-known functions decreasing sperm competition risk include manipulation of female propensity to re-mate and/or attractiveness (e.g. Chapman et al., 2003; LaFlamme, Ravi Ram, & Wolfner, 2012; Lung & Wolfner, 2001) and blocking her genital tract by forming plugs to prevent additional successful copulations, as commonly occurs in many taxa (Barker, 1994; Jia *et al.*, 2002; Mangels *et al.*, 2016; Sutter & Lindholm, 2016). On the other hand, the role of SFPs in sperm displacement has been reported for example in the fruit fly *Drosophila melanogaster* (Harshman & Prout, 1994) and in the seed beetle *Callosobruchus maculatus* (Yamane *et al.*, 2015), in which seminal fluid of the second male to mate with a female causes a reduction in the number of sperm from the previous mating and correlations have been found between allelic variation at SFP loci and levels of sperm displacement, as well as resisting ability to being displaced (Clark *et al.*, 1995; Fiumera *et al.*, 2005). Moreover, in polyandrous ants and bees, seminal fluid enhances the survival of own sperm, while preferentially eliminating sperm of rival males (Den Boer *et al.*, 2008, 2010).

So far, the literature on varying functions of seminal fluid modulating sperm competitive ability focuses mainly on separate-sexed organisms, but similar functions could also have evolved in hermaphrodites. Indeed, post-mating sexual selection has been suggested as a major evolutionary force shaping reproductive traits especially in simultaneous hermaphrodites (i.e. organisms with both male and female reproductive functions) (Charnov, 1979; Charnov & City, 1996; Michiels, 1998; Schärer *et al.*, 2015; Marie-Orleach *et al.*, 2016). Because frequent multiple mating is common in many reciprocally copulating simultaneous hermaphrodites, and individuals are capable of storing sperm from multiple ejaculate donors, there is an opportunity for selection to operate on differential fertilization success and resulting sperm competition among ejaculate donors (Baur, 1998; Michiels, 1998; Koene, 2005; Leonard, 2006; Anthes, 2010; Domínguez & Velando, 2013).

Our study organism, the flatworm *Macrostomum lignano*, is a reciprocally copulating simultaneous hermaphrodite in which self-fertilization does not occur (Schärer & Ladurner, 2003; Ladurner *et al.*, 2005b), and is an emerging model organism to study ejaculate adaptations driven by sperm competition and sexual conflict (e.g. Janicke & Schärer, 2009a; Schärer *et al.*, 2011; Marie-Orleach *et al.*, 2016; Patlar *et al.*, 2019, submitted). *M. lignano* can adjust its sex allocation, that is the strategic investment to produce eggs and sperm, in response to sperm competition level (e.g. Schärer & Ladurner, 2003; Janicke & Schärer, 2009a, 2010; Ramm *et al.*, 2019). It has been clearly shown that in larger social groups, where sperm competition intensity is high compared to small groups, worms invest more in their male sex function, as captured by traits such as testis size, testicular activity (Schärer *et al.*, 2004b), sperm production rate (Schärer & Vizoso, 2007) and spermatogenesis speed (Giannakara *et al.*, 2016). Moreover, it has been established that increasing investment in testis size and sperm production increases relative paternity success (Sekii *et al.*, 2013; Vellnow *et al.*, 2018). *M. lignano* also exhibits high mating rates, potentially due to the fact that individuals are motivated to donate sperm more than to receive it, just as in many other simultaneous hermaphrodites (Michiels, 1998; Schärer *et al.*, 2015; Greeff & Michiels, 2017). However, the high motivation to mate more, in general, also increases the risk of receiving (excess) sperm and/or seminal fluid, and concomitantly increases risks of polyspermy, sexually transmitted pathogens and/or receipt of manipulative SFPs (Charnov, 1979; Schärer *et al.*, 2015). Thus, it is likely that adaptations to gain control over the received ejaculate evolve in many simultaneous hermaphrodites, such as the sperm digestion common in gastropods (Baur, 1998; Michiels, 1998; Greeff & Michiels, 2017) and counter-adaptations to take control over own ejaculate such as bypassing the normal way of transferring sperm by hypodermic insemination in flatworms (Schärer *et al.*, 2011; Ramm *et al.*, 2015b; Ramm, 2016).

In *M. lignano*, there is a post-mating ‘suck behaviour’ that often occurs immediately after mating, which is proposed to be an adaptation to remove received ejaculate by sucking it out, that is potentially to gain control over the received ejaculate (Schärer *et al.*, 2004a, 2011; Vizoso *et al.*, 2010; Marie-Orleach *et al.*, 2013). Supporting this hypothesis about the function of suck behaviour, sperm have morphological adaptations to resist being removed by the recipient; each sperm possesses a frontal feeler and two stiff lateral bristles to anchor itself in the antrum (Schärer *et al.*, 2004a, 2011; Vizoso *et al.*, 2010). Moreover, a recently identified novel function of two seminal fluid transcripts, *suckless-1* and *suckless-2*, which potentially mediate sperm competitive ability in *M. lignano*, occurs not by directly influencing sperm interactions between rivals but instead by manipulating this suck behaviour of the partner and thereby affecting the fate of transferred ejaculates (Patlar *et al.*, submitted). We showed that the RNAi knock-down of these two transcripts in ejaculate donors increases the occurrence of the suck behaviour of their mating partner. This implies that the normal expression of these genes manipulates the partner to suck less often, thereby meaning more sperm can likely be retained in the partner’s sperm storage organ, potentially enhancing paternity success (Patlar *et al.*, 2019, submitted). We further showed substantial genetic variation in the expression of these transcripts (Patlar *et al.*, 2019), suggesting that this variation may be linked with variation in sperm competitive ability in *M. lignano*.

Although variation in sperm competitive ability can be expected to be depleted through strong directional selection, ejaculate traits often exhibit persistent genetic variation (Simmons & Kotiaho, 2002; Pitnick *et al.*, 2009; Simmons & Moore, 2009). One potential explanation for this paradox is the existence of genotype-by-environment interactions (GEIs) (Danielson-François *et al.*, 2006; Kokko & Heubel, 2008; Hunt & Hosken, 2014). GEIs create the potential to maintain genetic variation within populations exposed to conditions that vary in time and space (Via & Lande, 1985; Gillespie & Turelli, 1989), and could be especially important for understanding variation in sexually selected traits (Kokko & Heubel, 2008; Hunt & Hosken, 2014). So far, however, only a small number of studies have demonstrated GEI for sperm competitive ability itself (Engqvist, 2008; Lewis *et al.*, 2012), although several others have shown substantial GEI for sperm traits (Ward, 1998, 2000; Simmons & Kotiaho, 2002; Snook *et al.*, 2010; Nystrand *et al.*, 2011; Evans *et al.*, 2015; Marie-Orleach *et al.*, 2017) and one showed GEI for expression of seminal fluid transcripts (Patlar *et al.*, 2019), suggesting that variation through GEI can be widespread for ejaculate traits and potentially that it is related with, and could thereby help explain the maintenance of, variation in relative paternity success as an outcome of differential sperm competitive ability.

In this study, we therefore primarily aimed to investigate GEI for the expression of the *suckless-1* and *suckless-2* transcripts that potentially mediate sperm competition by manipulating partner suck behaviour, as well as for relative paternity success measured as first individual to mate (defensive sperm competitive ability; *P*_1_) or second individual to mate (offensive sperm competitive ability; *P*_2_) under sperm competition in *M. lignano*. We were then able to additionally test for a potential relationship between variation in seminal fluid expression generated through GEI and relative paternity success under sperm competition.

## Materials and methods

### Study organism

The free-living marine flatworm *Macrostomum lignano* (Schärer & Ladurner, 2003; Ladurner *et al.*, 2005b) cultures are kept in the laboratory at 20°C, 60% relative humidity, 14:10 light:dark cycle in 6-well tissue culture plates (Techno Plastic Products AG, Trasadingen, Switzerland) containing artificial sea water (ASW) with 32‰ salinity, and fed *ad libitum* with the diatom *Nitzschia curvilineata*. Under these conditions, worms frequently copulate and lay about 1-2 eggs per day (Schärer & Ladurner, 2003; Schärer *et al.*, 2004a). During reciprocal copulations, both individuals transfer sperm and seminal fluid to each other via the stylet (male copulatory organ), and received sperm are stored in their female antrum (sperm storage organ) (Schärer *et al.*, 2004a; Vizoso *et al.*, 2010). Seminal fluid is produced by prostate gland cells located around the stylet (Ladurner *et al.*, 2005a; Vizoso *et al.*, 2010), and includes a complex mixture of proteins (Weber *et al.*, 2018).

In this study, we conducted experiments to investigate GEI for relative paternity success and seminal fluid expression using four different inbred lines: DV8, DV13, DV28 and DV71 (Sekii *et al.*, 2013; Patlar *et al.*, 2019). The origin and maintenance of these inbred lines is explained elsewhere (Vellnow *et al.*, 2017; Patlar *et al.*, 2019). In a previous study, these chosen inbred lines (hereafter also referred to as *genotypes*) were found to be slightly different in their overall seminal fluid investment and their plastic response to different group size manipulations, suggested promising potential for GEI in seminal fluid transcript expression, but GEI was not clearly established, likely due to low statistical power (Patlar *et al.*, 2019). To asses sperm competitive ability of these chosen genotypes, we used another inbred line, DV1, which is the line commonly used in *M. lignano* studies (Janicke *et al.*, 2013), to generate standardized recipient worms and a Green-Fluorescent Protein (GFP)-expressing outbred culture to generate sperm competitors. Note that the GFP marker is a dominant allele expressed in all somatic and gametic cell types allowing us to easily and reliably genotype the offspring following a double mating trial between a wild type, non-GFP worm and a GFP-expressing worm in order to score paternity when in competition to fertilize the eggs of a wild-type non-GFP expressing recipient (Marie-Orleach *et al.*, 2014; Vellnow *et al.*, 2017). It has been shown that GFP-expressing worms are not affected by carrying the GFP marker in their reproductive traits compared to wild type outbred populations, which makes them reliable and powerful tools (Marie-Orleach *et al.*, 2014). The GFP expressing worms used in our experiment were from the outbred transgenic BAS1 culture (Marie-Orleach *et al.*, 2014; Vellnow *et al.*, 2017).

### Assessing sperm competitive ability of genotypes

Since the aim was to estimate GEI effects on seminal fluid expression and sperm competitive ability as a response to sperm competition level in their environment, we first raised genotypes in different social group sizes, namely pairs (group of two worms) and octets (group of eight worms). Following group size treatments, we assessed sperm competitive ability of these genotypes (focals) by conducting double mating trials in which virgin standardized mating partners (recipients) were mated sequentially with a focal worm followed by a (GFP-expressing) competitor, that is testing for the defensive sperm competitive ability of focals, *P*_1_; or else a competitor followed by a focal, that is their offensive sperm competitive ability, *P*_2_.

To manipulate social group size, we initially collected ca. 2-3 days old juveniles from main stock cultures of each genotype (ca. 150 per line – F_0_) and divided them into two glass Petri dishes with *ad libitum* food to let them grow and lay eggs. Once they started to reproduce, we collected their 2-3 day old offspring (F_1_) into one Petri dish (for randomization of juveniles collected from two Petri dishes of F_0_) and then we randomly distributed these F_1_ offspring into 24-well tissue culture plates, including 1 ml of ASW and *ad libitum* food in each well, to form groups of pairs and octets. Social groups were distributed on plates in a way that balanced for any potential plate position and genotype effects. In total, we formed 40 replicates (each comprising the eight different genotype/group size combinations) for the *P*_1_ assay and 41 replicates for the *P*_2_ assay). In order to avoid the potentially confounding effect of mismatched environmental conditions of competitors (Engqvist, 2013; Engqvist & Reinhold, 2016), we also raised GFP offspring either in pairs or octets (324 replicates each of pairs and octets) generated at the same time and under the same conditions as the focal genotypes, such that in each assay, the focal genotype and GFP competitor always matched in terms of prior social group size. Note that all the required offspring to form the social groups of genotypes and GFP competitors were collected within three days to minimise age differences. Thereafter, these offspring were raised for up to eight weeks in their given groups by transferring them to freshly prepared 24-well plates every week to prevent accumulation of their newly-hatched offspring.

In parallel, DV1 offspring needed for each double mating trial – to be used as unmated standardized recipients – were collected from the stock cultures (i.e. containing ca. 100 adult (F_0_) worms) and distributed individually on 24-well tissue culture plates (each well containing 1 ml ASW and *ad libitum* food) to keep them under strictly isolated conditions (in total, ca. 1300 isolated individuals). Approximately 24 hrs before each mating trial was conducted, we coloured recipients to distinguish them in mating pairs by transferring them individually into 60-well HLA Terasaki Plates (Greiner Bio-One, Frickenhausen, Germany), with each well containing 3 μl colour solution (5 mg of Colorant Alimentaire Grand Blue, Les Artistes, Paris, France per one ml 32‰ ASW) and 7 μl 32‰ ASW with food. Following the colouring step and before the mating trial itself, worms were briefly transferred to fresh 24-well plates without colour solution and food (including only 1ml of 32‰ ASW) for a few minutes for residual colouring to be washed out. In this way, worms were coloured slightly blue, which has no effect on worms’ maintenance, fecundity and mating behaviour (Marie-Orleach *et al.*, 2013), but which allows us to easily distinguish them from the focal worm under a stereomicroscope.

The double mating trials were initiated approximately eight weeks after the social group size manipulation of the focal and GFP competitors – and the isolation of recipients – began. In order to avoid pseudo-replication for both the focal and GFP competitors, one individual worm was picked randomly from each genotype/group size combinations and immediately used for the sperm competition assay for one assay only (and thereafter a subset of these was used for the gene expression measurements – see below). For logistical reasons, we divided the sperm competition assays into blocks performed over 13 days, with identical procedures on each day, and we ensured that recipients, genotypes and competitors used on each day were similarly aged (ranging from 55-62 days old) and randomly assigned.

Each day, we first paired a (unmated DV1) recipient (*recipient one*) for the first mating for two hours with a given individual (either a focal worm from one of the genotype/group size combinations or its GFP competitor, depending of the type of trial). At the end of this two hour period, the given individual was transferred to be immediately paired with a second recipient (*recipient two*) for a further two hours, while the recipient one was paired with the second individual to mate (either the GFP competitor or a focal worm from one of the genotype/group size combinations, opposite to the first period). Following this, we paired the second individual to mate with recipient two after removing the first individual. The aim of pairing each genotype and its competitor with two recipients sequentially was simply to increase the total number of offspring and thereby the precision of our paternity estimates, considering that individual worms lay relatively few eggs. Immediately after the total four-hour mating period, the focal worms were individually transferred to 1 ml tubes including 25 μl RNALater®, while GFP competitors were paired one-by-one with a separate group of virgin worms (DV1) in 24-well tissue plates, to verify their fertility. If a GFP worm did not sire any offspring with its partner after ca. seven-eight days, we paired it with at least three others to disentangle whether the GFP worm or its partner was the cause of the infertility. We later excluded data where GFP competitors did not achieve any reproductive success when paired with their additional partners.

Paternity assessment was done by counting the offspring of recipient one and two, which were isolated after the trials to let them lay eggs, by sorting GFP expressing or non-GFP expressing offspring under a stereomicroscope equipped with epi-fluorescence (Nikon SMZ-18 stereomicroscope with a Nikon C-HGFI Intensilight fluorescence and GFP filter cube; Nikon GmbH, Düsseldorf, Germany) over a period of two weeks after ensuring the last offspring were observed. In total, 234 out of 320 mating trials for the *P*_1_ assay and 248 mating trials out of 328 for the *P*_2_ assay were successfully measured for paternity success. The decrease in targeted sample size for *P*_1_ and *P*_2_ mating trials was due to lost worms during social group size treatment or due to excluded recipients if both (recipient one and two) did not produce offspring. In total, paternities for 2416 and 2291 offspring were assigned in the *P*_1_ and *P*_2_ assays, respectively.

### Seminal fluid transcript expression measurements

In order to measure the expression of *suckless-1* and *suckless-2*, we picked at random eight samples of each genotype/social group size combination from the worms used to assess *P*_1_. We performed RNA extraction of the *P*_1_ samples using the Reliaprep™ RNA tissue miniprep system (Promega, USA, #Z6112,) following the manufacturer’s instructions. Afterwards, reverse transcription was performed using 4.0 μl of 10.0 μl total RNA solution with the cDNA-Synthesis Kit H Plus (VWR Peqlab, Germany, 732-3273). CFX Connect™ Real-Time PCR Detection System (BioRad, CA, USA) was used for gene expression measurements. Reaction volumes were set at 10 μl, comprising 3 μl 2X SsoFast EvaGreen Supermix with low ROX (BioRad CA, USA), 150 nM of each primer pair (1.5 μl per pair), 3 μl nuclease-free water and 1 μl cDNA. Initial thermal cycling conditions were 1 cycle at 95°C for 5 minutes, followed by 39 cycles of denaturation at 95°C for 15 seconds and annealing/polymerization with a temperature of 59°C for 30 seconds. Raw Ct values (triplicated technical measurements for each sample and transcript) were extracted from CFX Software version 3.0. One sample, out of a total of 64, could not be quantified for gene expression because of possibly failed RNA extraction of this sample. We first evaluated the absolute expression values of technical replicates for consistency between them. To do so, we calculated the average value between all possible combinations of two replicates (average value of technical replicate one-two, one-three and two-three). If one technical replicate deviated by one Ct value or more from the other two (i.e. one Ct value increase is equal to ca. two times of mRNA/gene copies), we excluded this technical replicate (in total approximately 6% across all measurements) and based the expression measurement on the remaining two. Relative transcript expression values were calculated as ΔCt (Ct of gene of interest − Ct of internal control) after averaging technical replicates and results are reported as −ΔCt values (Schmittgen & Livak, 2008). The gene *macpiwi* (Pfister *et al.*, 2007) was used as internal control, which is a stable gene in terms of expression at different group sizes (Patlar *et al.*, 2019).

### Statistical analyses

All statistical analyses were performed with the R statistical software, version 3.2.3 (R Development Core Team, 2011). For transcript expression analyses, a two-way ANOVA approach was used to examine the main effects of genotype and social group size and the genotype-by-environment (group size) interaction, with relative transcript expression (−ΔCt) as the dependent variable. To examine the GEI for paternity success, we fitted generalized linear models with a binomial error distribution with logit link function (*P*_1_ and *P*_2_ analysed in separate models) and models to examine three-way interaction of genotype, group size and mating order (with the two recipients). All models for paternity success analyses included the response variable as a matrix where the first column is the number of focal and the second column is the number of GFP offspring, and a random effect of focal ID (due to the fact that each focal was measured for up to two recipients). Significance tests of interactions were based on Chi-square tests examining changes in deviance when the interaction term was dropped from the full model. Finally, we fitted general linear models with a binomial error distribution with logit link function to examine main effects of the expression of transcripts on *P*_1_ (i.e. the ratio of focal offspring to total offspring in the *P*_1_ assay), for recipient one and recipient two separately. The rationale of fitting separate models for recipient one and two was, first, to be able to evaluate gene expression effects on each recipient separately considering that transcript expression measurements were performed only following matings with recipient two, and second, because we had found an effect of the order of mating with the two recipients on *P*_1_ (see Results).

## Results

As predicted based on our previous results, seminal fluid transcript expression differs significantly between genotypes for both *suckless-1* and *suckless-2* and both exhibit significant GEI (Table 1). In addition, overall relative expression levels (−ΔCt) were similar between social groups (−3.79 ± 2.60 in pairs and −3.13 ± 2.65 in octets for *suckless-1* and 0.31 ± 1.83 in pairs and 0.73 ± 1.40 in octets for *suckless-2*). Although, the overall mean relative expression level of the two transcripts were apparently quite different (−3.46 for *suckless-1* and 0.52 for *suckless-2*), the reaction norms showing differential expression pattern of transcripts across social groups were strikingly similar for all genotypes (Fig. 1), suggesting GEI for expression of these transcripts manifests in a quite coordinated manner.

**Table 1:** Two-way ANOVA results for the seminal fluid transcript expression variation. The significant *P*-values are written bold. (SS: Sum of squares).

| Effect | df | *suckless-1* |  |  | *suckless-2* |  |  |
| --- | --- | --- | --- | --- | --- | --- | --- |
|  |  | SS | *F* | *P* | SS | *F* | *P* |
| Group size | 1 | 6.20 | 1.87 | 0.18 | 2.63 | 1.31 | 0.26 |
| Genotype | 3 | 132.01 | 13.24 | **<0.001** | 30.30 | 5.02 | **<0.01** |
| Interaction | 3 | 87.58 | 8.78 | **<0.001** | 18.98 | 3.15 | **0.03** |
| Residuals |  | 166.20 |  |  | 106.56 |  |  |

**Figure 1:**
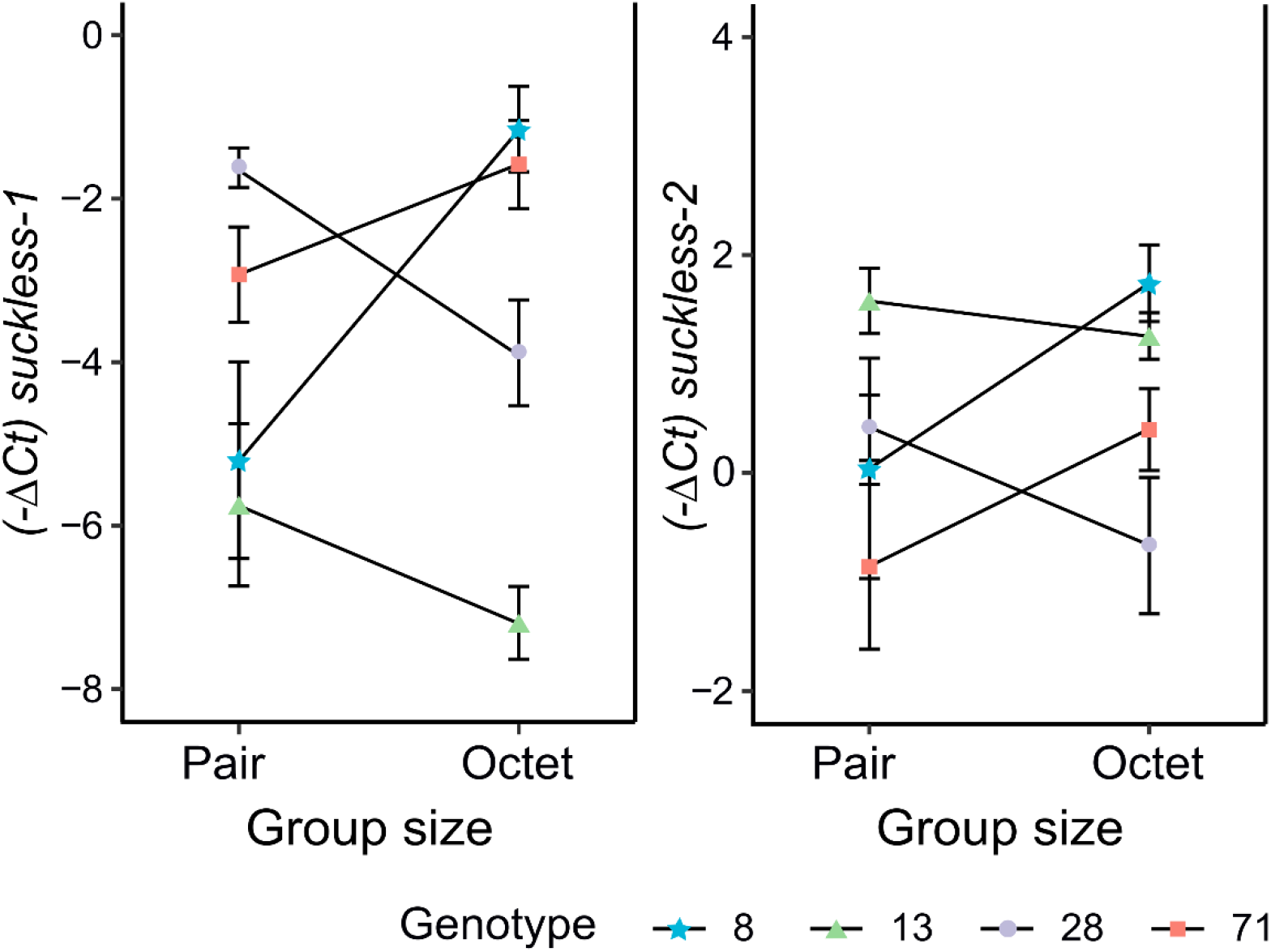
Seminal fluid gene expression of genotypes across social group size treatments. The mean relative expression (−ΔCt) of *suckless-1* (left) and *suckless-2* (right) of genotypes (inbred lines DV8, DV13, DV28 and DV71) in pairs and octets (social group sizes). Error bars show the standard errors of the mean.

For the sperm competition assay, the initial model comparisons including mating order with recipient one (or two) as a discrete factor, plus genotype, social group size and their two-way interaction, showed that paternity success (scored as *P*_1_ and *P*_2_) does not exhibit GEI (Table 2). However, based on the different reaction norms of genotypes observed for recipient one and two (Fig. 2), and a significant main effect of mating order shown in this model at least for *P*_2_ (and a similar, marginally non-significant trend for *P*_1_) (Table S1), we (retrospectively) preferred to analyse these data by instead fitting a model including a three-way interaction between genotype, social group size and mating order with the recipient. This model indeed shows that there is a significant three-way interaction, meaning genotypes differ in their relative paternity success depending on both the social group size and mating order (Table 2 & Table S2). Note that when we fitted models for GEI for each recipient separately, GEI was highly significant for both recipients in the *P*_1_ assay (*P* = 0.002 for recipient one, *P* = 0.005 for recipient two) and for the first recipient in the *P*_2_ assay (*P* < 0.001 for recipient one, *P* = 0.17 for recipient two) (Table S3).

**Table 2:** Model comparisons to evaluate interaction effects for paternity success. Generalized linear model comparisons (*P*_1_ and *P*_2_ as in separate models) based on likelihood ratio tests to evaluate the effects of two-way interaction (genotype-by-group size interaction) and three-way interaction (genotype-by-group size-by-mating order with the recipients). The full model for two-way interaction comparison incudes mating order, genotype and group size plus genotype-by-group size interaction as fixed factors, and focal ID as a random factor. The full model for three-way interaction comparison incudes mating order, genotype and group size plus all possible interactions as fixed factors, and focal ID as a random factor. The outcomes of the full models were added as supplementary tables (Table S1&S2).

| | $P_1$ | | | $P_2$ | | |
| --- | --- | --- | --- | --- | --- | --- |
| | Df | Chi | $P(>Chi)$ | Df | Chi | $P(>Chi)$ |
| **Model comparison for two-way interactions** |  |  |  |  |  |  |
| **Model (full)** | 10 | 3.99 | 0.26 | 10 | 6.51 | 0.09 |
| **Model without two-way interaction** | 7 |  |  | 7 |  |  |
| **Model comparison for three-way interactions** |  |  |  |  |  |  |
| **Model (full)** | 17 | 11.15 | **0.01** | 17 | 10.48 | **0.02** |
| **Model without three-way interaction** | 14 |  |  | 14 |  |  |

**Figure 2:**
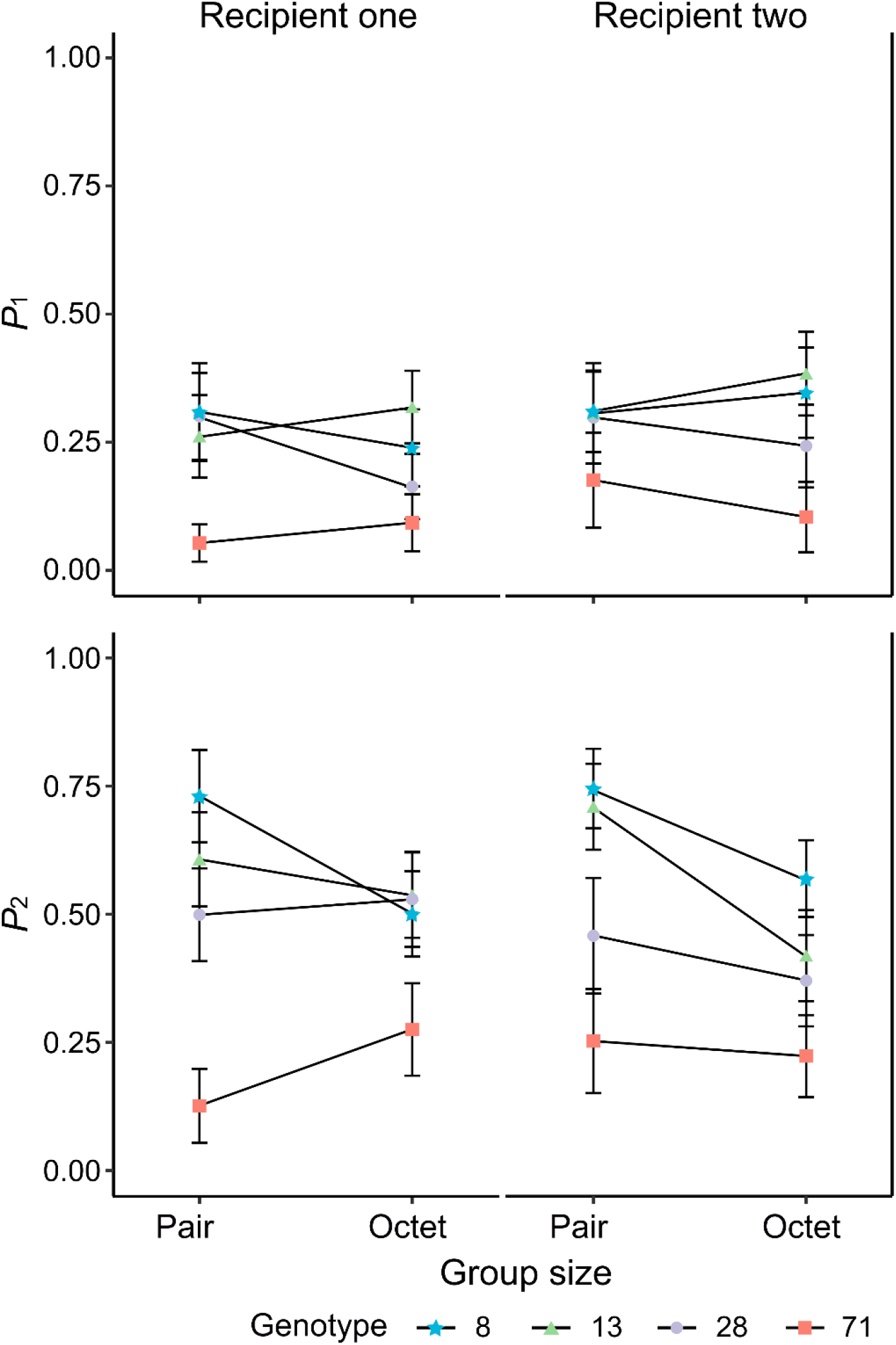
Relative paternity success (*P*_1_ and *P*_2_) of genotypes across social group size treatments in recipient one and two. The mean *P*_1_ and *P*_2_ scores of different genotypes (inbred lines) across group size treatments were shown for recipient one (left) and recipient two (right). *P*_1_ and *P*_2_ was scored as the ratio of offspring number sired by focal to the total offspring number produced by each recipient, representing paternity success as first or second to mate with recipients, respectively.

Finally, to determine how variation in seminal fluid transcript expression generated through GEIs might be associated with variation in paternity success, we regressed seminal fluid transcript expression data of individuals with their *P*_1_ scores. According to the generalized linear model fitted including additive main effects of *suckless-1* and *suckless-2*, only the expression of *suckless-1* had an effect on paternity success, but this was highly significant and consistent across both recipient one (*P* =0.006) and recipient two (*P* < 0.001) (Table S4). We also calculated the correlation between the expression of the two genes, which was marginally significant but weak (*r* = 0.27, *P* = 0.037). Based on this, we then dropped *suckless-2* from the models and evaluated and visualized only the effect of *suckless-1* (Figure 3) for each recipient (GLM for recipient one: *z* = 3.14, *P* < 0.001, recipient two: *z* = 3.96, *P* < 0.001). To be precise, the average coefficient for *suckless-1* = 0.24 (slope of log-odds for recipient one = 0.24398 and recipient two = 0.2302), which is interpreted as the expected change in log odds for a one-unit increase in the expression level of *suckless-1*. The odds ratio can be calculated by exponentiating this value to get 1.27 which means we expect to see about a 27% increase in the odds of paternity success of genotypes overall, for a one-unit increase the expression level of *suckless-1*.

**Figure 3:**
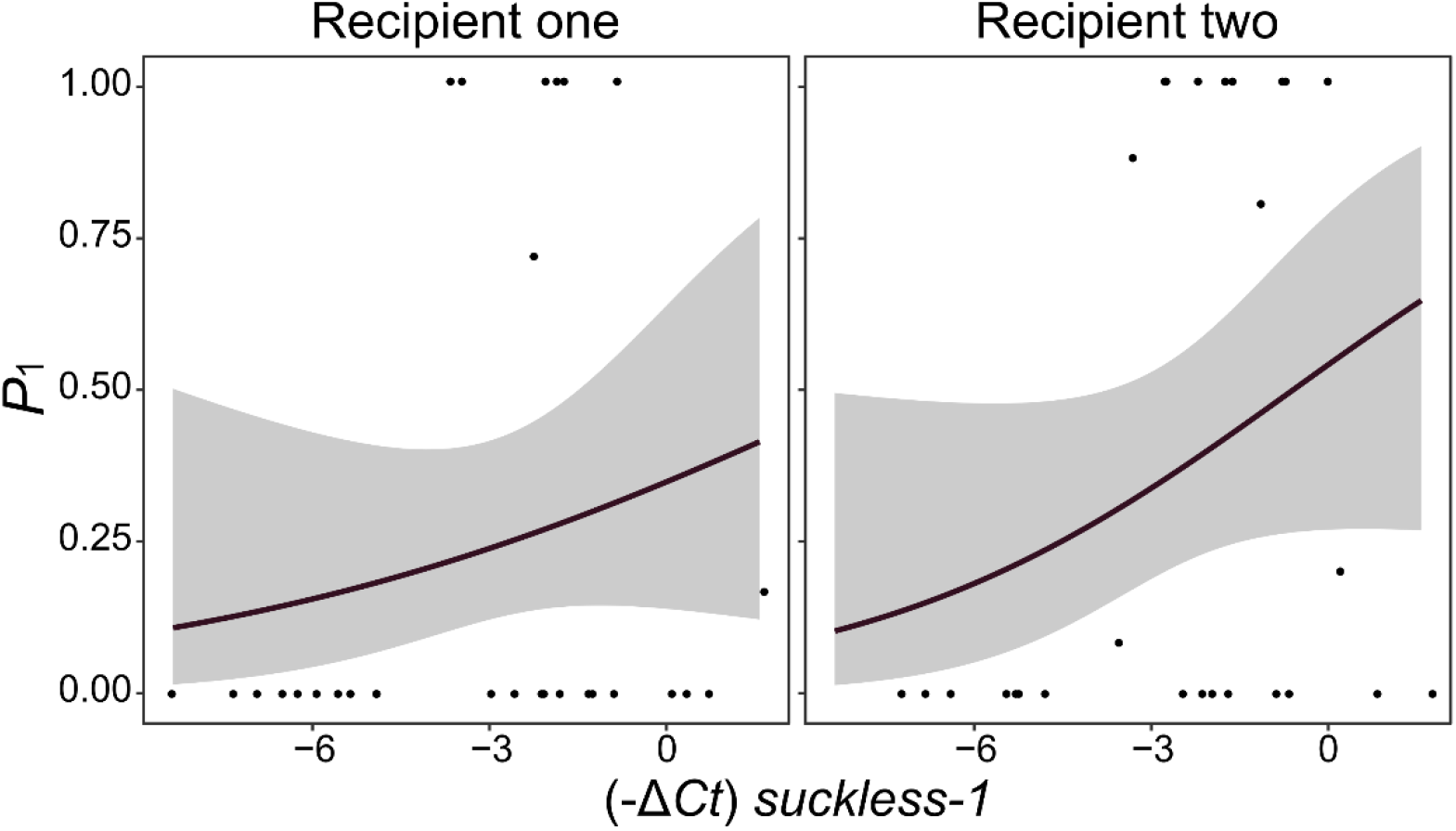
Predicted relative paternity success (*P*_1_) depending on relative expression of *suckless-1* in recipient one and two. Curves were drawn using a binomial logistic regression model; the shaded areas indicate 95% confidence intervals for each curve.

## Discussion

We found evidence of GEI for the expression level of two seminal fluid transcripts that were recently functionally characterised, *suckless-1* and *suckless-2* (Patlar *et al.*, submitted) and for relative paternity success, as well as evidence linking *suckless-1* expression to sperm competitive ability. Research on GEI in sexually selected traits is an important area of study in evolutionary biology, and has been proposed to explain the maintenance of standing genetic variation in sexually selected traits that are often under strong selection (Pomiankowski & Moller, 1995; Clark, 2002; Kokko & Heubel, 2008; Ingleby *et al.*, 2010; Hunt & Hosken, 2014). The majority of studies have focused on either sexually selected traits involved in the pre-mating episode of sexual selection (reviewed by Ingleby et al. 2010) or for sperm traits that were proposed as possible adaptations to sperm competition (reviewed by Reinhardt et al. 2015). These studies have shown GEI on sperm characteristics such as sperm length (Morrow *et al.*, 2008), sperm transfer rate (Engqvist, 2008), sperm velocity (Evans *et al.*, 2015) and sperm mobility (Purchase *et al.*, 2010), whereas others have focused on testis size as a predictor of sperm production (Nystrand *et al.*, 2011; Marie-Orleach *et al.*, 2017). To our knowledge, there is only one previous study that directly tested GEIs for seminal fluid transcript expression, our own previous investigation also in *M. lignano* (Patlar *et al.*, 2019; but see also Mangels *et al.*, 2015). Here, we therefore provide novel evidence of GEI for another major aspect of the male ejaculate of adaptive significance under sperm competition.

First of all, we supported our previous results regarding significant genotypic variation for the relative expression of *suckless-1* and *suckless-2*, beyond which we extended our findings by demonstrating also significant GEI for these transcripts. We already had some evidence of GEI for seminal fluid transcript expression in our previous study where we showed significant GEI for 14 out of 58 seminal fluid transcripts (Patlar *et al.*, 2019), though this did not include *suckless-1* or *suckless-2*, and where we also showed a lack of significant social group size effect for all 58 transcripts (Patlar *et al.*, 2019; but see Ramm *et al.*, 2019). In contrast to several studies that found high degrees of differences in expression of SFPs between manipulated sperm competition levels, the lack of group size, and thus sperm competition effect in *M. lignano* may therefore be explained by the existence of GEI. Overall, these results suggest that GEI could be widespread among seminal fluid production and this potentially can help explain the maintenance of standing genetic variation for seminal fluid proteins that were shown in some studies (for gene expression: Smith *et al.*, 2009; Patlar *et al.*, 2019; for protein abundance: Baer *et al.*, 2012; Goenaga *et al.*, 2015; Mangels *et al.*, 2015). In fact, one can expect some genetic variation to be maintained for some SFPs because of the lack of fitness relation, thus selection do not act on but, considering their varying functions related tightly with successful reproduction, this can be unlikely a reason for standing genetic variation (Heifetz *et al.*, 2001; LaFlamme *et al.*, 2012; Schjenken & Robertson, 2014; Avila *et al.*, 2015, see also Poiani, 2006), this is unlikely to be a general explanation for genetic variation in SFPs. Yet phenotypic studies that have manipulated social environments to manipulate sperm competition have revealed considerable plasticity for seminal fluid expression according to the presence/absence of rival males (e.g. Harris & Moore, 2005; Fedorka *et al.*, 2011; Ramm *et al.*, 2015; Mohorianu *et al.*, 2017; Sloan *et al.*, 2018). Here, we argue that variation in social environment and thus the extent of GEIs may arise from heterogeneity in social conditions and could maintain genetic variation in seminal fluid expression. Further studies are needed to explain how often GEI occurs among organisms for seminal fluid expression, and another interesting question will be to what extent seminal fluid gene expression is controlled by genetic/epigenetic mechanisms to imprint social environmental heterogeneity to gene expression (Perry & Mank, 2014).

Second, we found some evidence for GEI for both *P*_1_ and *P*_2_, revealing that GEIs occur for relative paternity success and implying genotype-by-environment interaction effects on donor sperm competitive ability. Notably, however, the differing reaction norms of different genotypes depended not just on the social group size, but also on mating order (i.e. whether we assessed paternity success in matings with the first or second recipient) in *M. lignano*. To the best of our knowledge, only a few studies have so far examined GEI for sperm competitive ability, and one study provides very similar evidence to ours of GEI for *P*_1_, but in that case depending on nutritional conditions affecting ejaculate contents in the flour beetle *Tribolium castaneum* (Lewis *et al.*, 2012). Another study showed GEI for sperm transfer rate also depending on nutritional conditions, indirectly suggesting a GEI effect on paternity success depending on the amount of sperm transferred in the scorpionfly *Panorpa cognata* (Engqvist, 2008). Other studies mainly focused on genotype-by-*genotype* interactions for sperm competitive ability in other organisms, provided strong evidence for female-male interactions and male-male interactions explaining variation in paternity success measured as *P*_1_ and/or *P*_2_ (Clark *et al.*, 1999, 2000; Bjork *et al.*, 2007; Dowling *et al.*, 2007; Firman, 2014).

Both GEI and individual genotype interactions for sperm competitive ability may well be taxonomically widespread, since many factors affect paternity success in sperm competition. For example, our results also indicate that the pattern of GEI depended crucially on the mating order. Our experimental design involved a focal worm being taken out from its respective group (pair or octet) and at once paired with a virgin recipient (one) for two hours, and thereafter immediately with a second virgin recipient (two) in the subsequent two hours. It has been shown that group size manipulation affects mating rate and average sperm transfer rate in *M. lignano* (Janicke & Schärer, 2009a; b), and worms raised in pairs have more stored sperm in their seminal vesicle than worms raised in octets as a result of high mating rate in octets (Janicke & Schärer, 2009b; Marie-Orleach *et al.*, 2014). It is possible that only few sperm are left in the seminal vesicles in worms raised in octets, and the genotypes which were used in this study may vary considerably among themselves for sexual traits. Therefore, genotypes might well differ in their testis size, sperm production rate or the amount of sperm stored in their seminal vesicles, and this may differentially affect ejaculate size and composition transferred to first *versus* second recipients – and thereby relative paternity success – depending on the group size and genotype. For example, after mating with recipient one, some genotypes might be faster or slower to replenish and restore sperm and/or seminal fluid proteins when paired with the second recipient, or genotypes might differ in their motivation to re-mate with novel partners depending on the group size from which they originated.

A further interesting finding from our study is that variation in expression level of *suckless-1* generated through GEIs robustly predicted defensive sperm competitive ability (*P*_1_), across both first and second recipients, potentially suggesting adaptive GEI for seminal fluid gene expression. Note that we did not evaluate offensive sperm competitive ability (*P*_2_) in terms of its relation with seminal fluid expression because sperm displacement occurs in *M. lignano* (Marie-Orleach *et al.*, 2014) and it has been proposed that the shape of the copulatory organ could be important in sperm competition to outcompete sperm of the previous donor by mediating sperm displacement (Janicke & Schärer, 2009a): we therefore considered that paternity measurements of *P*_2_ assays might also depend strongly on variation in stylet morphology of the chosen genotypes. Moreover, when the worm mates first a previously unmated recipient then only its sperm/seminal fluid affects whether the partner sucks or not, whereas if it mates second then potentially both its and the first donor’s sperm/seminal fluid can affect recipient behaviour, confounding our test.

We expected that an increase in the expression of seminal fluid transcripts, assuming their expression is positively correlated with the amount of protein transferred, should result in an increased number of sperm being retained in the recipient’s sperm storage organ by manipulating the suck behaviour. Therefore, relative paternity success may be linked with the expression of these transcripts, and indeed that appears to be the case for one of them, *suckless-1*. However, our conclusion about the underlying mechanism should be treated with caution, because we did not directly manipulate transcript expression and it is likely that several other seminal fluid transcripts, as well as the other aspect of male allocation, sperm production, will to some extent co-vary with *suckless-1* expression (Patlar *et al.*, 2019). Indeed, we previously demonstrated positive genetic correlation among seminal fluid transcripts and between testis size and seminal fluid investment (Patlar *et al.*, submitted), and recent studies clearly showed that increases in testis size have a positive impact on paternity success in *M. lignano* (Sekii *et al.*, 2013; Vellnow *et al.*, 2018). Therefore, the effect we found might be due to the collinearity between sperm and seminal fluid production, or between different seminal fluid components. If more than just *suckless-1* is involved in such a response, this could help explain why the natural variation in *suckless-1* expression investigated here was clearly linked to relative paternity success, but in a previous manipulative experiment where we knocked down the expression of (only) *suckless-1* we observed no such clear impact on paternity success (Patlar *et al.*, submitted). In the curent study, it was striking that *P*_1_ of some recipients (from both recipient one and two groups) scored as one hundred percent (see Fig. 3), particularly linked with higher expression level of *suckless-1* suggesting that an increase in seminal fluid expression, and especially *suckless-1* or transcripts that are highly positively genetically correlated in their expression with *suckless-1* (Patlar *et al.*, 2019), might have an antagonistic effect on re-mating rate of the partner. Nevertheless, we note that we did not see any evidence for a link between paternity success and the expression level of a second seminal fluid gene, *suckless-2*. Further studies are needed to test for the actual effect of *suckless-1* on paternity success in competitive environments, especially by controlling the other aspects of male allocation. In fact, there are promising tools such as manipulation of sperm production in a dose-dependent manner in *M. lignano* (Sekii *et al.*, 2013) that could be a very useful approach to control for the effect of variation in sperm production/transfer and thereby help disentangle the independent effects of seminal fluid proteins on sperm competitive ability.

To conclude, our study demonstrates that GEI occurs for seminal fluid transcript expression, depending on the social group size, and thus the level of sperm competition, and additionally GEI also occurs for sperm competitive ability but depending on the both group size and potentially on other traits e.g. mating rate or average sperm transfer rate which themselves also depend on the level of sperm competition and individual genotypes. Further, we found evidence that natural variation in expression level of *suckless-1* through GEIs could predict relative paternity success, and so influences the outcome of sperm competition.

## Acknowledgments

We thank A. Giannakara and M. Weber for help maintaining the laboratory cultures and T. Schmoll for useful statistical feedback. This study was funded by the Deutsche Forschungsgemeinschaft (DFG, grant number RA 2468/1-1 to SAR).

## Author contributions

BP and SAR developed the experimental design to assess relative paternity success. BP performed the experiments; collected the data; and performed data analyses. BP wrote the manuscript with the contribution of SAR.

