## Supplemental Tables for "Genotype-by-environment interactions for seminal fluid expression and sperm competitive ability"

**Table S1: Results of binomial generalized models testing the effect of group size manipulation, genotype, mating order and genotype-by-group size interaction on paternity success, $P_1$ and $P_2$.** Abbreviations represent social group size (S), genotype (G), and mating order with recipients (R). The significant *P*-values are written bold.

| | $P_1$ | | | | $P_2$ | | | |
|---|---|---|---|---|---|---|---|---|
| **Effect** | **Estimate** | **SE** | ***z*** | ***P*** | **Estimate** | **SE** | ***z*** | ***P*** |
| **Intercept** | -1.55 | 0.70 | -2.12 | **0.03** | 2.96 | 0.86 | 3.44 | **<0.001** |
| **R(two)** | 0.30 | 0.17 | 1.80 | 0.07 | -0.47 | 0.18 | -2.68 | **<0.01** |
| **G(DV13)** | -0.48 | 0.92 | -0.52 | 0.60 | -1.21 | 1.16 | -1.05 | 0.30 |
| **G(DV28)** | -0.19 | 0.93 | -0.21 | 0.84 | -3.07 | 1.15 | -2.68 | **<0.01** |
| **G(DV71)** | -3.01 | 1.10 | -2.75 | **<0.01** | -6.78 | 1.35 | -5.03 | **<0.001** |
| **S(octet)** | -0.51 | 0.93 | -0.55 | 0.59 | -2.68 | 1.09 | -2.46 | **<0.01** |
| **G(DV13):S(octet)** | 1.21 | 1.24 | 0.98 | 0.33 | 0.69 | 1.50 | 0.46 | 0.64 |
| **G(DV28):S(octet)** | -1.28 | 1.28 | -0.99 | 0.32 | 2.36 | 1.53 | 1.55 | 0.21 |
| **G(DV71):S(octet)** | -0.54 | 1.45 | -0.37 | 0.71 | 3.80 | 1.67 | 2.27 | **0.02** |
| **Random effect** | **Variance** | | **Obs.** | **Groups** | **Variance** | | **Obs.** | **Groups** |
| **Focal ID** | 8.44 | | 351 | 234 | 13.13 | | 369 | 248 |

**Table S2: Results of binomial generalized models testing the effect of group size manipulation, genotype, mating order and all possible interactions on paternity success, $P_1$ and $P_2$.** Abbreviations represent (social) group size (S), genotype (G), and mating order with recipients (R). The significant *P*-values are written bold.

| | $P_1$ | | | | $P_2$ | | | |
|---|---|---|---|---|---|---|---|---|
| **Effect** | **Estimate** | **SE** | ***z*** | ***P*** | **Estimate** | **SE** | ***z*** | ***P*** |
| **Intercept** | -1.61 | 0.74 | -2.18 | **0.03** | 2.57 | 0.95 | 2.71 | **<0.01** |
| **R(two)** | 0.64 | 0.58 | 1.10 | 0.27 | 0.01 | 0.52 | 0.03 | 0.98 |
| **G(DV13)** | -0.20 | 0.96 | -0.21 | 0.84 | -1.06 | 1.27 | -0.84 | 0.40 |
| **G(DV28)** | 0.34 | 0.98 | 0.35 | 0.73 | -2.84 | 1.27 | -2.24 | **0.03** |
| **G(DV71)** | -3.52 | 1.34 | -2.63 | **0.01** | -7.12 | 1.48 | -4.81 | **<0.001** |
| **S(octet)** | -0.39 | 0.97 | -0.40 | 0.69 | -2.61 | 1.20 | -2.18 | **0.03** |
| **R(two):G(DV13)** | -0.80 | 0.68 | -1.18 | 0.24 | 0.16 | 0.72 | 0.22 | 0.83 |
| **R(two):G(DV28)** | -1.49 | 0.74 | -2.01 | **0.05** | -0.19 | 0.86 | -0.22 | 0.82 |
| **R(two):G(DV71)** | 0.15 | 1.02 | 0.15 | 0.88 | 0.39 | 0.82 | 0.47 | 0.64 |
| **R(two):S(octet)** | -0.49 | 0.69 | -0.72 | 0.47 | 0.28 | 0.66 | 0.43 | 0.67 |
| **G(DV13):S(octet)** | 0.39 | 1.28 | 0.30 | 0.76 | 0.99 | 1.64 | 0.61 | 0.55 |
| **G(DV28):S(octet)** | -2.16 | 1.35 | -1.60 | 0.11 | 2.77 | 1.68 | 1.66 | 0.10 |
| **G(DV71):s(octet)** | 0.47 | 1.85 | 0.26 | 0.80 | 5.37 | 1.86 | 2.89 | **<0.001** |
| **R(two):G(DV1):S(octet)** | 1.94 | 0.85 | 2.27 | **0.02** | -1.44 | 0.93 | -1.54 | 0.12 |
| **R(two):G(DV28:S(octet)** | 2.23 | 1.00 | 2.22 | **0.03** | -1.29 | 1.05 | -1.23 | 0.22 |
| **R(two):G(DV71):S(octet)** | -1.07 | 1.64 | -0.65 | 0.52 | -3.76 | 1.26 | -2.97 | **<0.001** |
| **Random effect** | **Variance** | | **Obs.** | **Groups** | **Variance** | | **Obs.** | **Groups** |
| **Donor ID** | 8.55 | | 351 | 234 | 13.13 | | 369 | 248 |

**Table S3: Model comparisons to evaluate genotype-by-environment interaction (GEI) effects for paternity success across recipients.** Generalized linear model comparisons (*$P_1$* and *$P_2$* for recipient one and recipient two as in separate models) based on likelihood ratio tests to evaluate the effects of two-way interaction (genotype-by-group size interaction). The full model for two-way interaction comparison incudes genotype and group size plus genotype-by-group size interaction as fixed factors, and focal ID as a random factor.

| | ***$P_1$*** | | | ***$P_2$*** | | |
|---|---|---|---|---|---|---|
| | **Df** | **Deviance** | ***P*(>Chi)** | **Df** | **Devience** | ***P*(>Chi)** |
| **Model comparison for GEI - Recipient one** | | | | | | |
| **Model (full)** | 178 | -15.05 | **0.002** | 185 | -23.42 | **<0.001** |
| **Model without GEI** | 175 | | | 182 | | |
| **Model comparison for GEI - Recipient two** | | | | | | |
| **Model (full)** | 163 | -12.78 | **0.005** | 174 | -5.02 | 0.17 |
| **Model without GEI** | 160 | | | 171 | | |

**Table S4: Models to evaluate effects of suckless-a and suckless-2 on paternity success measured as defensive sperm competitive ability (*$P_1$*) across recipients.** Generalized linear models were fitted including main effects of expression level of *suckless-1* and *suckless-2*, and focal ID as a random factor on a response variable as a matrix where the first column is the number of focal and the second column is the number of GFP expressing offspring.

| | Recipient one | | | | Recipient two | | | |
|---|---|---|---|---|---|---|---|---|
| **Effect** | **Estimate** | **SE** | ***z*** | ***P*** | **Estimate** | **SE** | ***z*** | ***P*** |
| **Intercept** | -0.82 | 0.37 | -2.21 | **0.03** | -0.19 | 0.27 | -0.71 | 0.48 |
| ***suckless-1*** | 0.25 | 0.09 | 2.74 | **0.006** | 0.23 | 0.06 | 3.62 | **<0.001** |
| ***suckless-2*** | -0.07 | 0.14 | -0.51 | 0.61 | -0.06 | 0.10 | -0.54 | 0.59 |
